# Disassociating cerebral vasomotion from low frequency spontaneous neurovascular coupling

**DOI:** 10.1101/2025.06.23.661067

**Authors:** Runchong Wang, Priya Patel, Luke Boorman, Michael Okun, Clare Howarth, Jason Berwick

**Affiliations:** School of Psychology, University of Sheffield, Sheffield, UK; Opteran, Sheffield, UK; School of Life Sciences, University of Nottingham, Nottingham, UK; Neuroscience Institute, University of Sheffield, Sheffield, UK

**Author notes:** These authors contributed equally to this work (Runchong Wang, Priya Patel, Jason Berwick).

## Abstract

Vasomotion, vascular oscillations at ∼0.1 Hz, may serve as a biomarker and therapeutic target for neurodegenerative diseases, but its origins, structure across brain vasculature, and correlation with neural activity remain unclear. This study examined the spatiotemporal characteristics of cerebral vasomotion and its relationship to neural activity in anaesthetised Hooded Lister rats using simultaneous recordings of neuronal activity and haemodynamics in motor and whisker barrel cortices. In a subset of rats, tissue oxygen was also measured. Blood pressure was pharmacologically modulated to alter vascular oscillations. We found that vasomotion was driven by the arterial tree. Two prominent activity patterns emerged: global vasomotion across the entire hemisphere and phasic vasomotion seen as a travelling wave running through the surface arteries. Moreover, vasomotion was associated with low tissue oxygen and was largely independent of spontaneous neural activity and therefore not a product of neurovascular coupling.

## Introduction

Functional magnetic resonance imaging (fMRI) has transformed the way we understand the workings of the brain, allowing scientists to take measurements from the entire brain in both active human and animal subjects^1^. The technique also has great potential for the development of functional biomarkers of disease. However, there is a fundamental issue at play, namely that fMRI does not measure neuronal activity but changes in cerebral haemodynamics. These haemodynamic changes are driven by a mechanism called neurovascular coupling (NVC).

fMRI for both sensory evoked and spontaneous activity^2,3^ relies on the tacit assumption that the fMRI Blood Oxygen Level Dependent (BOLD) signal, produced by NVC mechanisms, is stable, constant and unaffected by disease conditions. However, it is increasingly apparent that dysfunction of the neurovascular unit and therefore NVC is an important co-morbidity of many brain disorders, may speed-up disease progression and even can be responsible for starting the disease in the first place^4^. A potential candidate as an early biomarker of neurovascular breakdown is a blood flow oscillation called vasomotion.

Vasomotion is a pervasive low frequency oscillation that can occur in any tissue in the body at a frequency of ∼0.1Hz (one cycle every 10 seconds). Its existence has been known for decades, indeed its first appearance can be traced back to the original Roy and Sherrington paper in 1890, which has often been cited as the first paper to propose the existence of NVC^5^. Vasomotion, especially in animal research, has often been associated with a poor preparation rendering the experiment useless^6^. However, little research has been done investigating why vasomotion occurs, its vascular structure within the brain and whether it is independent of neuronal activation or in some way interacts with ongoing baseline neuronal activity. One of the main ideas is that vasomotion exists to ensure that a compromised microcirculation can obtain enough oxygen for vulnerable tissue^7^. With the emergence of whole brain neurovascular imaging in the awake mouse the term vasomotion was coopted to refer to spontaneous NVC oscillations occurring at low frequencies^8,9^, whereas the original interpretation to a large extent has been lost^10,11^. Vasomotion has the potential to be both an important biomarker of early brain pathology and a therapeutic method to boost brain blood flow in neurodegenerative diseases. Toward these ends, we need to understand cerebral vasomotion and how it interacts with baseline neuronal activity.

Here, we manipulated baseline blood pressure to elevate or inhibit distinct vasomotion oscillations in urethane-anaesthetised rats. Neuronal, haemodynamic activity and tissue oxygenation changes were measured simultaneously. We show that vasomotion oscillations occur at frequencies close to 0.1Hz (rather than a broad spectrum of frequencies as reported in mouse studies), are driven by the arterial tree, and are associated with low tissue oxygen. We show that vasomotion, at least in the rat, is distinct from baseline low frequency spontaneous NVC.

## Results

In the beginning of each experiment, we localised the barrel and motor cortices by measuring haemodynamic responses with 2-dimensional optical imaging spectroscopy (2D-OIS, Fig. 1 and Methods) following 2s whisker stimulation. We then inserted silicon probes into both the whisker and motor regions (Fig. 1d and Methods) and proceeded with continuous 2D-OIS and electrophysiology data collection over a period of 2100s. Example recording data is shown in Fig 2. An important effect of urethane anaesthesia is that it lowers blood pressure. Therefore, in all recordings, starting at 290s, phenylephrine was continuously infused, which gradually restored blood pressure to normal physiological levels (90-110mmHg, Fig. 2g). In a representative animal a clear oscillation of the blood volume signal (Hbt) can be seen in the low blood pressure period which dissipates later (Fig. 2h). Power spectrum of Hbt in this initial period shows a dominant peak close to 0.1Hz which is the frequency of cerebral vasomotion (Fig. 2i). This animal also exhibited spontaneous burst suppression neuronal activity, another known property of urethane anaesthesia^12^. Prior to phenylephrine infusion the bursts of neuronal activity were not always associated with Hbt elevation (Fig. 2j). However, after the removal of vasomotion each burst of neuronal activity results in a subsequent peak in Hbt (Fig. 2k-l; see below for more detailed analysis).

**Figure 1:**
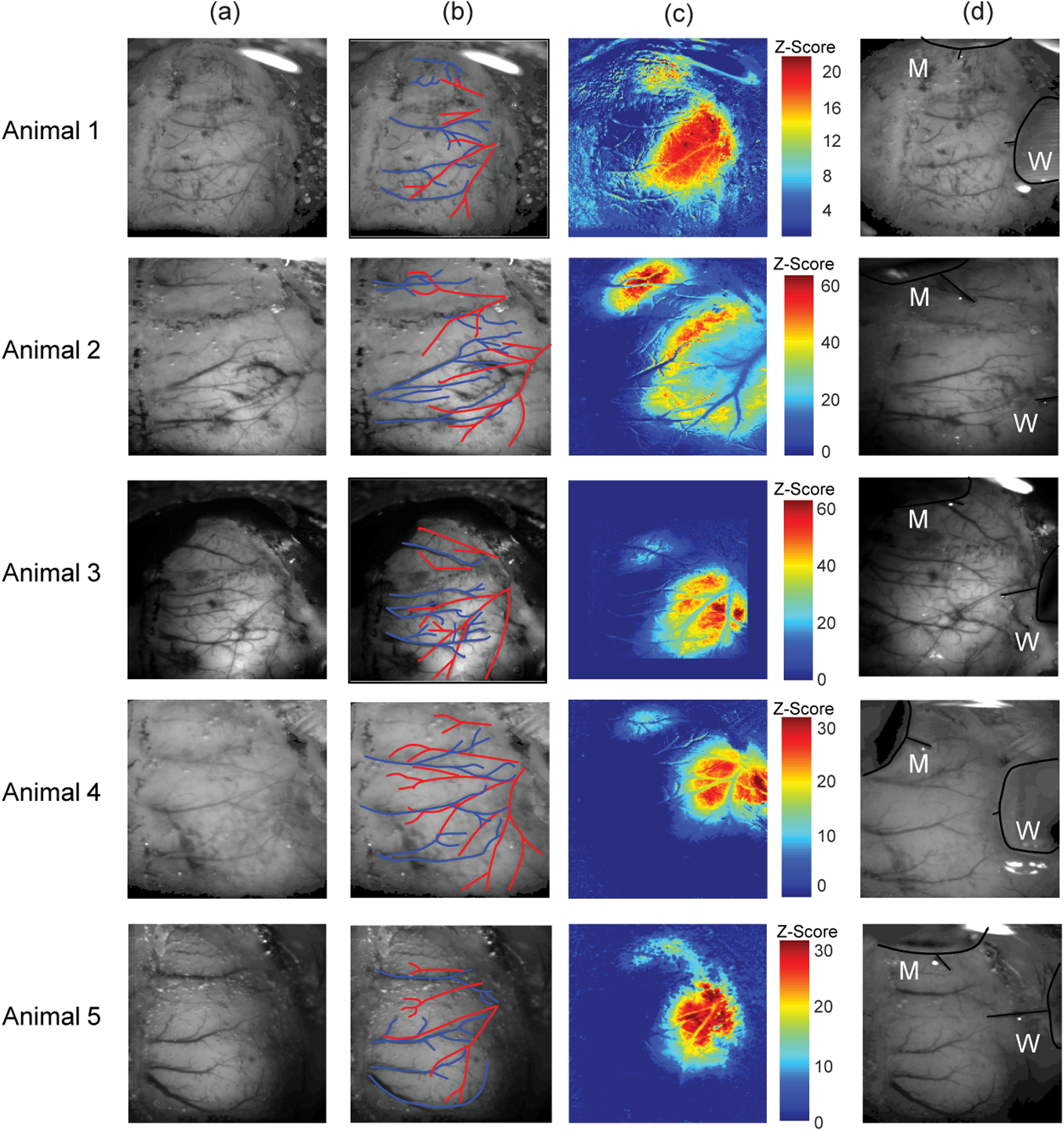
Experimental preparation and location of electrode insertion from 5 animals. **a)** In vivo grey level images of the thinned cranial window preparation over the right cerebral hemisphere. **b)** Overlay of the major arteries (red) and veins (blue) running over the surface of the cerebral cortex. **c)** Activation blood volume maps to a 2s left whisker pad stimulation following General Linear Model analysis. Red areas represent pixels that most correspond to a representative hemodynamic response function. Two response areas emerge which reflect the larger whisker barrel cortex (lateral and posterior in the window) and the motor cortex (more medial and anterior). **d)** Same image as shown in a) but with both the whisker (W) and motor (M) cortex electrode inserted into the brain to enable simultaneous neurovascular coupling measurements.

**Figure 2:**
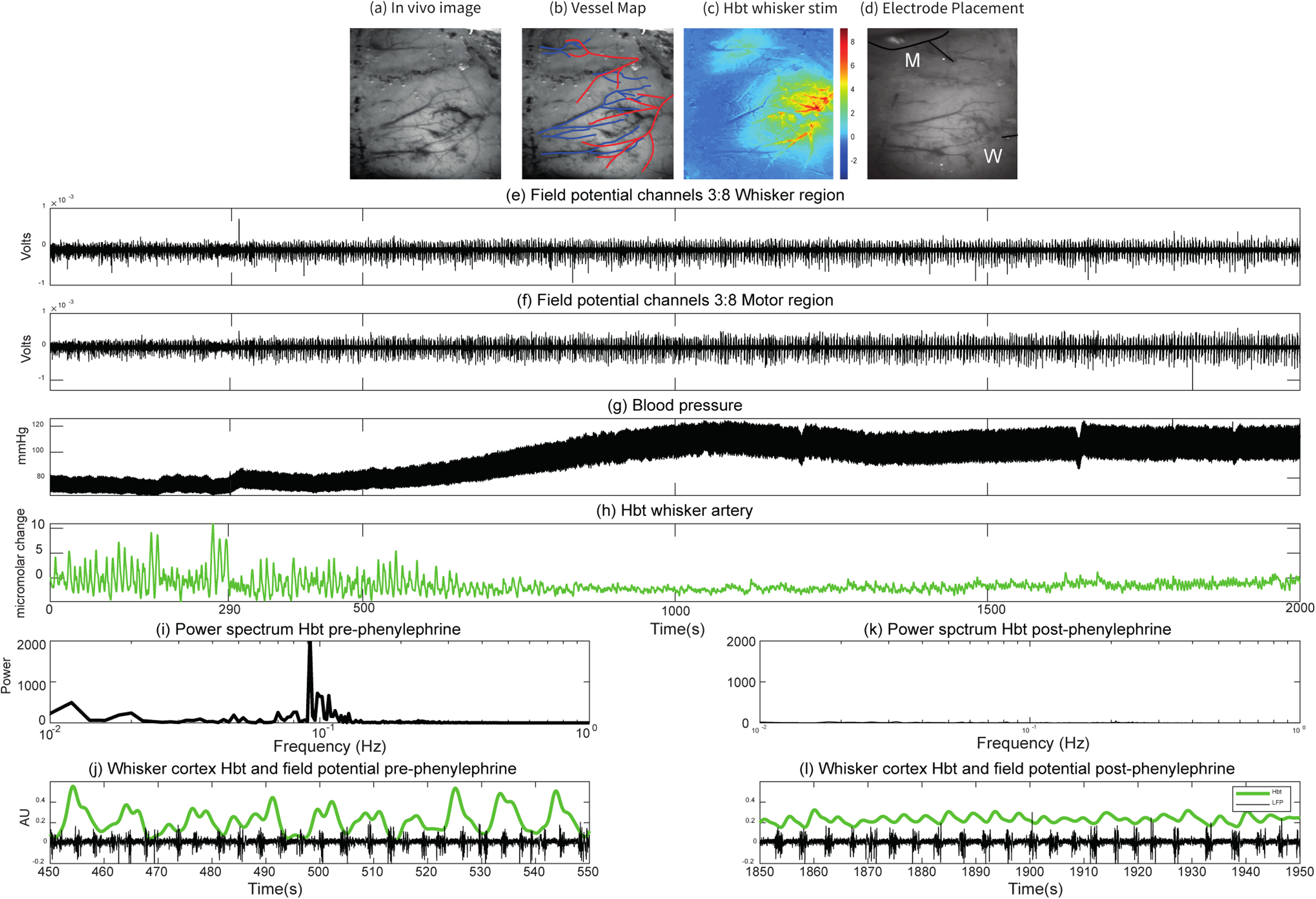
Representative responses from one animal. Phenylephrine infusion was started after 290s. a) In vivo grey level image of cortical surface. **b)** Overlay of the major arteries (red) and veins (blue) running over the surface of the cerebral cortex. **c)** Activation blood volume maps to a 2s left whisker pad stimulation following General Linear Model analysis. **d)** Image showing whisker (W) and motor (M) cortex electrode inserted into the brain to enable simultaneous neurovascular coupling measurements. **e)** Average local field potential (LFP) response from electrode channels 3 to 8 from the whisker barrel cortex electrode. **f)** Average LFP response from electrode channels 3 to 8 from the motor cortex electrode. **g)** Blood pressure measurement. **h)** Total haemoglobin (Hbt) from the whisker barrel artery. **i)** Power spectrum of the Hbt time series from 450:550 seconds showing a strong vasomotion peak at 0.1Hz. **j)** Zoomed in time series of Hbt (green) and whisker cortex field potential (black) time series between 450:550 seconds. The LFP time series shows marked burst suppression. **k)** Power spectrum of the Hbt time series from 1850:1950 seconds. The peak in vasomotion has disappeared. **l)** Zoomed in time series of Hbt (green) and whisker cortex field potential (black) time series between 1850:1950 seconds. The LFP time series shows burst suppression, which is reflected in the Hbt time series.

### Low blood pressure enhances cerebral vasomotion and reduces brain oxygenation

There was a significant increase in mean arterial blood pressure (MABP) after phenylephrine injection (*M* = 90.5mmHg, *SD* = 13.8) compared to before phenylephrine injection (*M* = 69.9 mmHg, *SD* = 10.4, *t*(16) = -12.314, p<0.001).

The change in MABP produced a marked effect on the power of vasomotion. We measured vascular responses from 24 main branches of the Middle Cerebral artery (MCA), of which 18 were from the artery overlying the whisker barrel cortex and the remaining 6 were overlying the motor cortex. In several animals, there was no identifiable artery over the motor cortex so the analysis could not be performed. For each branch, Hbt signal was measured. Power spectra were estimated from the 0-375s interval that represented the low blood pressure period, and from the 1400-1775s interval where MABP had normal physiological levels (Fig. 3a-c). The magnitude of vasomotion was taken as power in the 0.08-0.15Hz interval. For the whisker region artery (single sample taken from each rat), the power of vasomotion (Hbt) was higher (*Mdn* = 5731) before phenylephrine injection than after (*Mdn* = 326). A Wilcoxon Test indicated that this was statistically significant, (W = 0.00, z = -3.724, *p*<0.001). For the motor region artery (from 6 animals), the power of vasomotion (Hbt) was higher (*Mdn* = 2268) before phenylephrine injection than after (*Mdn* = 330). A Wilcoxon Test indicated that this was statistically significant, (W = 1.00, z = -1.992, *p* = 0.046).

**Figure 3:**
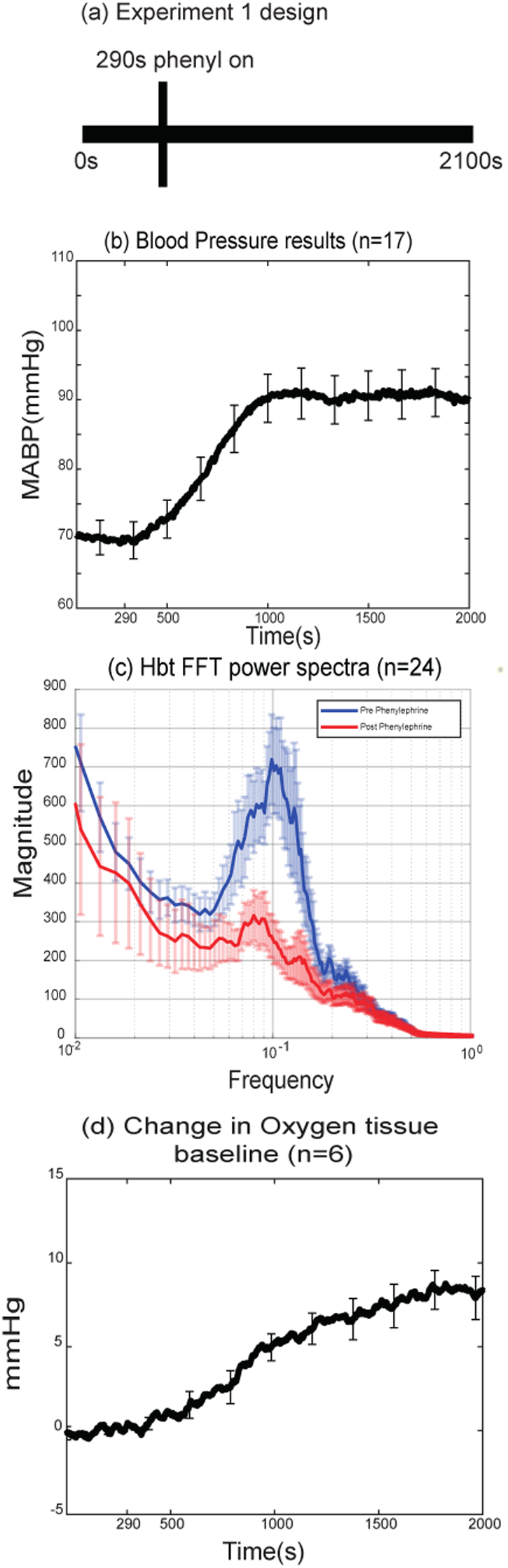
Blood pressure, vasomotion and tissue oxygen changes resulting from phenylephrine infusion. **a)** Schematic showing the 2100s experiments where continuous hemodynamic and neural data were collected. Phenylephrine infusion began at 290s. **b)** Averaged blood pressure change measured in 17 animals. **c)** Power spectrum of Hbt before (blue) and after (red) phenylephrine infusion. **d)** Change in tissue oxygen baseline (in millimetres of mercury, mmHg) in six animals following phenylephrine infusion at 290s. Dara = mean, error bars = standard error of the mean.

The increase in MABP and the reduction in vasomotion were also associated with an increase in baseline brain tissue oxygenation. In the 6 animals where tissue oxygen levels were measured with an oxygen electrode (see Methods) we observed a significant increase in oxygen tension in the cerebral cortex after phenylephrine injection (*M* = 19.3 mmHg, *SD* = 5.5) compared to before phenylephrine injection (*M* = 11.7 mmHg, *SD* = 2.8), *t*(5) = -6.493, p = 0.001, Fig. 3d). Taken together these results show that there was enhanced cerebral vasomotion in urethane-anaesthetised rats, which was associated with low oxygen tension in the cerebral cortex, which was reduced following blood pressure stabilization.

### Spatial relationship of vasomotion across the cortical surface

To investigate vasomotion dynamics, we identified the times of Hbt peaks and averaged the signal in the 20 seconds surrounding each peak (see Methods). Data from two example animals is shown, the first (Figure 4) reveals demonstrates vasomotion as a prominent vascular wave spreading across the cortex aligned with the direction of the main branches of the MCA. The second example (Fig. 5a-f) demonstrates vasomotion that was of a more global nature, emerging at all branches of the MCA at a similar time. To calculate the temporal lag of vasomotion between the branches of the artery, we used the most proximal and distal regions of the MCA, which were furthest apart. Across all animals the lag in Hbt between proximal and distal branches of the MCA was 0.95s ±0.18s (*SEM*) with the proximal branch leading (paired t-test, p=0.00046). Therefore, in general vasomotion can be seen as an oscillation close to 0.1 Hz that rapidly spreads across the cortex aligned to the vascular territories of the MCA.

**Figure 4:**
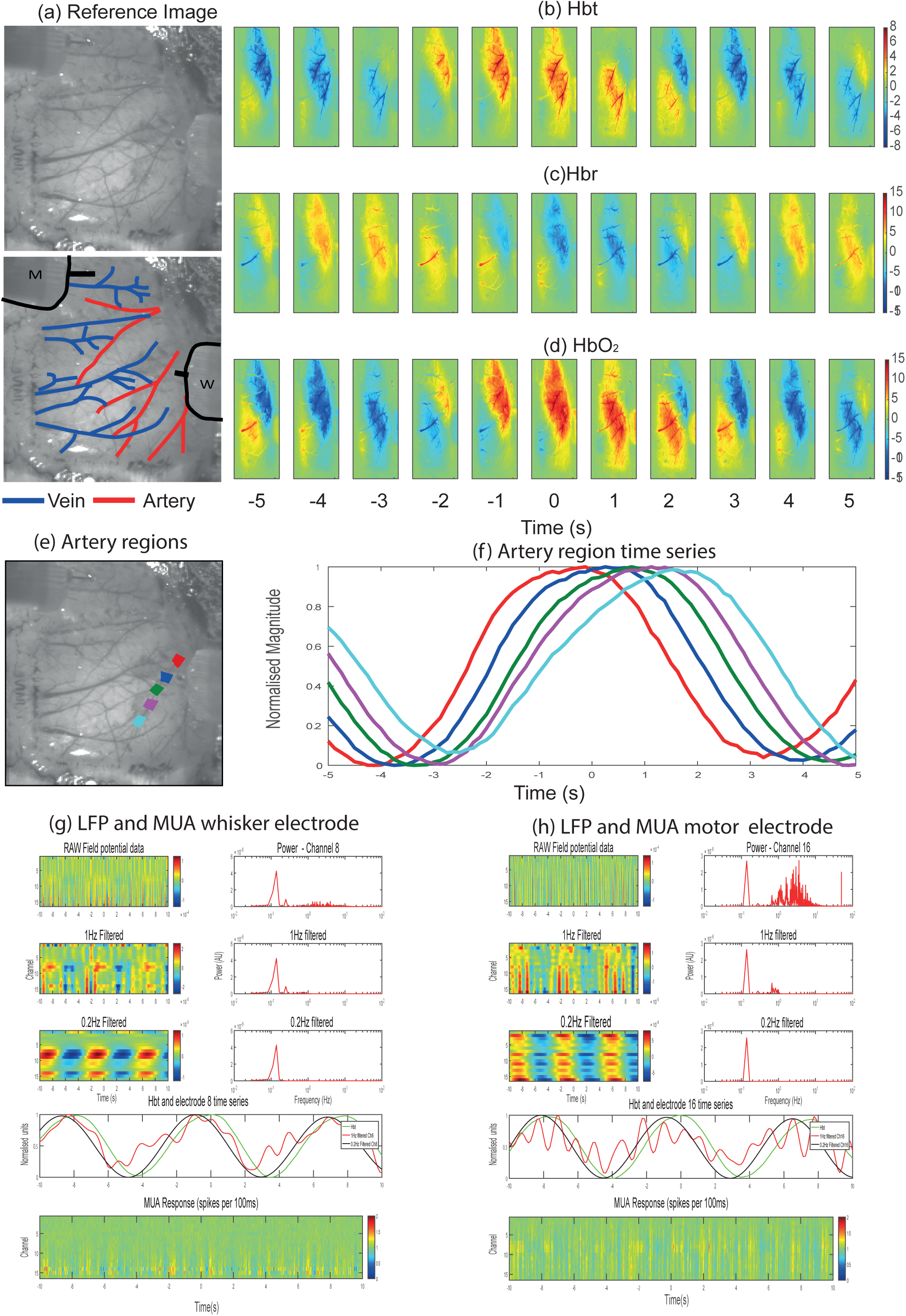
Representative animal showing phasic vasomotion. **a)** In vivo reference images (top) with (bottom) highlighted arteries (red), veins (blue) and electrode insertion locations (black, M = motor, W = whisker). **b)** Averaged spatial montage of total haemoglobin, Hbt with the peak of vasomotion aligned at time zero. Each image represents the average of 1s (8 frames) of data. Vasomotion can be seen as a band moving throughout the cortex aligned with the geometry of the middle cerebral artery (MCA). Colour bar represent ΔHbt. **c)** Averaged spatial montage of deoxyhaemoglobin, Hbr, with the peak of vasomotion aligned at time zero. Hbr is more aligned to the geometry of the main draining veins towards the midline of the brain. Colour bar represent ΔHbr. **d)** Averaged spatial montage of oxyhaemoglobin HbO_2_ with the peak of vasomotion aligned at time zero. This follows a similar pattern as the Hbt data. Colour bar represent ΔHbO_2._ **e)** In vivo image in which 5 regions of one branch of the MCA have been selected from proximal (closest to the heart, red) to distal (light blue). **f)** Hbt time series from the 5 selected regions. In this animal, there is a distinct phase lag along the vessel length of ∼2s. **g)** LFP and Multi-Unit Activity (MUA) from whisker region electrode extracted from the same time as the peak Hbt from vasomotion trials. Top row shows the raw LFP as a function of depth over time with a corresponding power from channel 8 of the electrode. A distinct peak at 0.1Hz can be seen. Second and third rows are from the same data but filtered below 1Hz and 0.2Hz respectively. The fourth row shows the normalised Hbt time series along with the LFP time series from channel 8 filtered at 1 and 0.2Hz. The hemodynamic signal lags the neuronal signals by less than one second. The bottom row shows the MUA data across all neural channels, little of no 0.1Hz oscillation is present. **h)** LFP and MUA from motor region electrode (channel 16) extracted from the same time as the peak Hbt from vasomotion trials, similar to **g)**.

**Figure 5:**
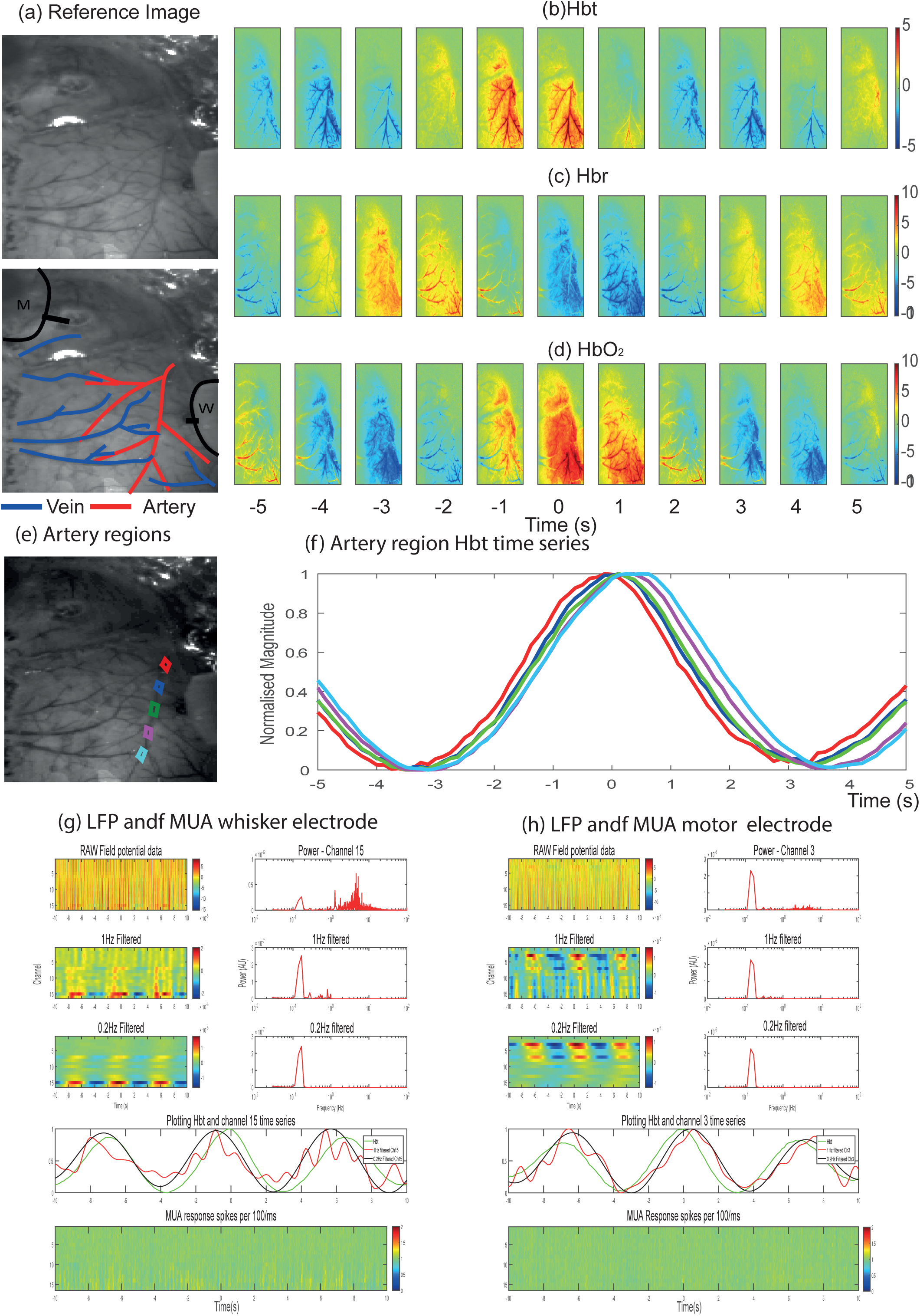
Representative animal showing global vasomotion. **a)** In vivo reference images (top) with (bottom) highlighted arteries (red), veins (blue) and electrode insertion locations (black, M = motor, W = whisker). **b)** Averaged spatial montage of total haemoglobin, Hbt, with the peak of vasomotion aligned at time zero. Each image represents the average of 1s (8 frames) of data. Vasomotion emerges from MCA across its entire length at the same time. Colour bar represent ΔHbt. **c)** Averaged spatial montage of Hbr with the peak of vasomotion aligned at time zero. Colour bar represent ΔHbr. **d)** Averaged spatial montage of HbO_2_ with the peak of vasomotion aligned at time zero. This follows a similar pattern as the Hbt data. Colour bar represent ΔHbO_2._ **e)** In vivo image where 5 regions of one branch of the MCA have been selected from proximal (closest to the heart) to distal. **f)** Hbt time series from the 5 selected regions. In this animal, there all sections increase close to the same time. **g)** LFP and MUA from whisker electrode extracted from the same time as the peak Hbt from vasomotion trials. Top row shows the raw LFP as a function of depth over time with a corresponding power from channel 15 of the electrode revealing a distinct peak at 0.1Hz. Second and third rows are from the same data but filtered below 1Hz and 0.2Hz respectively. The fourth row shows the normalised Hbt time series along with the LFP time series from channel 15 filtered at 1 and 0.2Hz. The hemodynamic signal lags the neuronal signals by less than one second. The bottom row shows the MUA data across all neural channels, little or no 0.1Hz oscillation present. **h)** LFP and MUA from motor region electrode (channel 3) extracted from the same time as the peak Hbt from vasomotion trials, similar to **g)**.

### Does neural activity drive cerebral vasomotion?

To understand if vasomotion correlates with neural activity, we examined the simultaneously recorded LFP and MUA data. In the aforementioned two representative animals, the LFP data contained an oscillation at the same frequency as the Hbt signal on several of the probe channels, but this varied as a function of depth across both the whisker and motor cortices (Fig. 4g-h, Fig. 5g-h). The MUA data (bottom panels of Fig. 4g-h, Fig. 5g-h) showed no notable oscillation around 0.1Hz.

To isolate the relationship between Hbt and the LFP signal we extracted a single neural channel from 11 probe sites (from 8 animals where both whisker and motor electrodes were used in 3 animals) where there was strong vasomotion and little or no burst suppression activity (burst suppression was a larger neural signal that dominated over the smaller 0.1Hz LFP signal). The average of this shows a clear oscillation in LFP response. Overall, the neural signal led the Hbt oscillation by 0.43 ± 0.04s *SEM* (Fig. 6a). To understand how this corresponds to spontaneous NVC, we extracted burst suppression haemodynamic peaks in the absence of strong vasomotion after blood pressure stabilisation (Fig. 6b). These animals showed clear responses in both LFP and MUA prior to the peak in Hbt. The lag between the MUA peak and Hbt was 2.11± 0.05s *SEM*. A final comparison was made between resting state (Fig. 6a and b) and 2s whisker evoked stimulation, and as expected both the LFP and MUA responses preceded the Hbt peak response with the peak Hbt response lagging 3.9± 0.07s *SEM* (Fig. 6c). Taken together these results show that spontaneous NVC driven by neuronal burst suppression and stimulation evoked NVC responses have the expected structure, where neuronal activity in both LFP and MUA data drive subsequent haemodynamics. For vasomotion we did see an associated LFP, but this was closely aligned in terms of phase, with little or no MUA neural correlate. Therefore, it can be concluded that vasomotion around 0.1Hz is not a product of normal NVC relationships.

**Figure 6:**
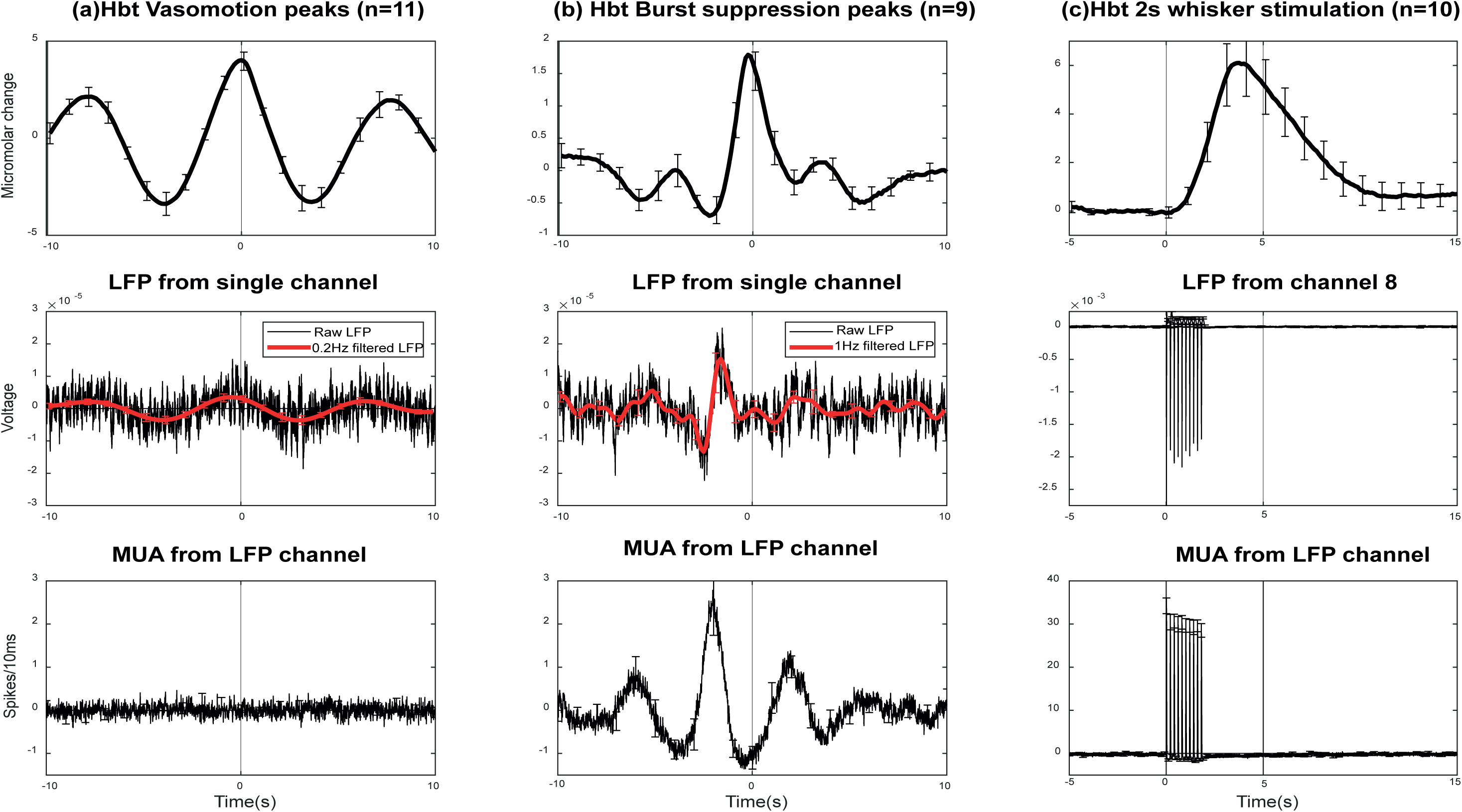
Comparing the phase relationship between vasomotion, spontaneous burst suppression, and stimulation induced neurovascular responses. **a)** Vasomotion from 11 sessions (from 8 animals, 3 used both whisker and motor electrodes), where there was strong vasomotion with little or no burst suppression. Top row averaged Hbt trials aligning the Hbt peak at time zero. Middle row LFP selected from a single channel in the same experiment, with a clear vasomotion peak at 0.1Hz. Bottom row MUA from the same LFP channel with no oscillation present. **b)** Same responses as described in a) but from animals that showed strong neuronal burst suppression in the absence of vasomotion in whisker cortex. The peak of Hbt (top row) seen at is preceded by both LFP (middle row) and MUA (bottom row) responses occurring ∼2s before the peak of Hbt. **c)** Responses from 2s whisker stimulation. The peak of Hbt responses occurs ∼4s after the onset of LFP and MUA neural activity. This represents the phase difference expected to be seen from stimulation induced neurovascular coupling. Data = mean, error bars = Standard error of the mean.

### Kernel analysis

Finally, we used the spontaneous neural data to assess how well it could predict the subsequent haemodynamics with and without the presence of vasomotion. Linear kernel linking Hbt to LFP signals was computed for each recording, along with the correlation between Hbt predicted from LFP using this kernel and the actual Hbt signal. The kernel captures the features of NVC. Three representative animals are shown in Figure 7. The first animal had vasomotion with no burst suppression, the correlation between measured and predicted Hbt was 0.15, which is close to noise level (Fig. 7a). The second animal had strong burst suppression with little or no vasomotion and Hbt prediction was much better, with a correlation coefficient of 0.62 (Fig. 7b). The third animal provides an example of whisker stimulation epoch, and here it can be seen that the predicted Hbt replicated the measured Hbt with a correlation of 0.9 (Fig. 7c).

**Figure 7:**
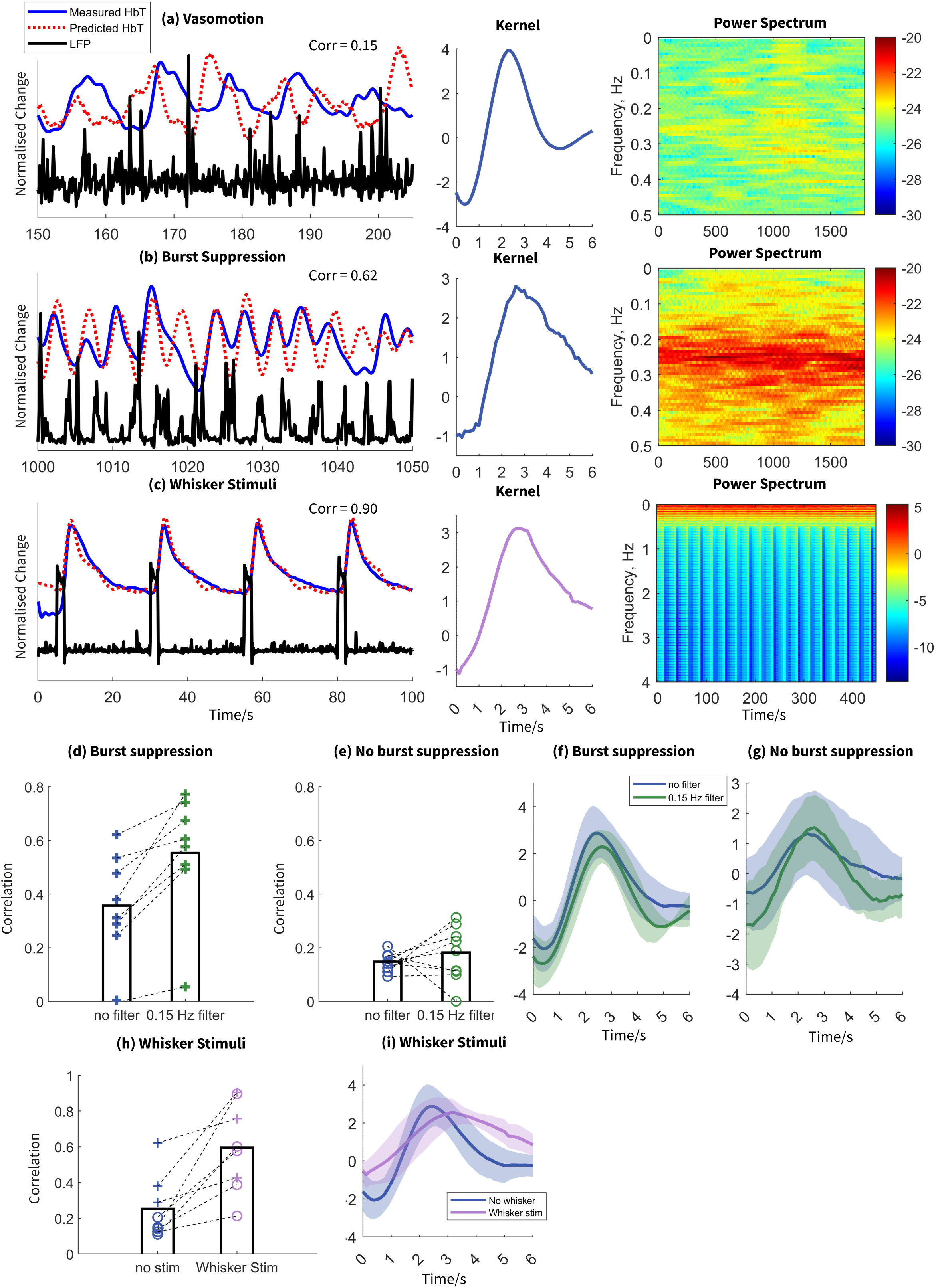
Burst suppression influences the relationship between vasomotion and NVC. Example subjects illustrating vasomotion without burst suppression **a)**, with vasomotion and burst suppression **b)**, and under whisker stimulation **c)**. The black line represents gamma-band LFP, the blue line represents measured Hbt, and the red dashed line represents predicted Hbt. The middle column is the kernel for the three subjects. The blue line indicates kernel for resting state and the purple line indicates kernel for whisker stimulation. The right column is power spectra of the LFP. **d)** and **e)**: Correlation between predicted and measured Hbt in subjects with and without burst suppression. The dark blue and green indicates data without and with 0.15 Hz filter, respectively. The cross and circle indicates subjects with and without burst suppression, respectively. **f)** and **g)**: Average kernel between gamma-band LFP and Hbt in subjects with **(f)** and without **(g)** burst suppression. The shaded region indicates standard error of the mean. **h)** Correlation between predicted and measured Hbt in subjects with (purple) and without (blue) whisker stimulation. The cross and circle indicates subjects with and without burst suppression, respectively. **i)** Average kernel between gamma-band LFP and Hbt in subjects with (purple) and without (blue) whisker stimulation. The shaded region indicates standard error of the mean.

In animals with strong burst suppression and vasomotion we assessed whether removal of vasomotion by filtering it out from the Hbt time series would improve the predictive power of the kernel analysis. In 8 sessions, a high-pass filter (0.15 Hz) was used to remove vasomotion from the Hbt signal. This increased the correlation coefficient from 0.33 to 0.50 (*z* = -2.70, *p* = 0.01, Fig. 7d). In animals with no burst suppression and vasomotion the Hbt prediction did not improve with 0.1Hz haemodynamic filtering (Fig. 7e). This suggests that either these animals’ NVC was dysfunctional or within the spontaneous anaesthetised condition the neural activity was low and did not drive detectable spontaneous NVC relationships. To test this, we analysed the predictive Hbt response in animals subjected to 2 seconds of whisker stimulation (*N* = 8, including 3 animals exhibiting burst suppression). The Hbt response was significantly elevated in the whisker stimulation condition compared to the no burst suppression condition (mean whisker = 0.53, mean no burst suppression = 0.14, *z* = -2.02, *p* = 0.04, Figure 7h). Therefore, NVC was preserved in these animals in response to a stimulus, indicating the relationships seen in the spontaneous condition were likely a result of neural activity being below a threshold that evoked vascular responses.

## Discussion

In this study, we have characterised cerebral vasomotion in the anaesthetised rat preparation. We have shown that, when present, vasomotion exhibits a large distinctive peak at 0.1Hz driven by changes in Hbt in the middle cerebral artery. It is not driven by coherent neural activity, but when present without burst-suppression has an LFP neural correlate occurring within 0.5s of the hemodynamic oscillation. Vasomotion was also associated with low tissue oxygen levels. In terms of spontaneous NVC responses if 0.1Hz was filtered out of the hemodynamic data the kernel prediction method became more accurate (but only when animals had robust burst-suppression spontaneous neural activity). This further supports the hypothesis that classically defined vasomotion is separate and distinct from spontaneous NVC relationships that spontaneous functional connectivity research relies upon.

### Vasomotion in rat compared to mouse

The results presented here are in line with in rat experiments over several years. Hudetz’s studies characterised the precise nature of the oscillation in rat sensory cortex^10,11^. Vasomotion has also been observed in many peripheral tissues^13–15^ again supporting the hypothesis that it is independent of, and not driven by, neuronal activity. We have confirmed and extended these studies with a more rigorous spatial analysis and with simultaneously acquired multi-channel neuronal activity. However, over the past decade, with the emergence of wide-field mouse imaging studies^16^ the term vasomotion has been used to describe spontaneous NVC connectivity results. Mateo et al ^8^ were the first to use this definition, elegantly showing that the power envelope of GAMMA band neural activity was coupled to ‘vasomotor’ hemodynamic oscillations around the broad peak of 0.1Hz, this effectively re-defined what vasomotion was and potentially has caused confusion for the field.

A major focus of research has now moved onto understanding vasomotion in mouse models of disease^9,17,18^. This has gained further importance with the suggestion that vasomotion might increase the clearance of toxic proteins seen in neurodegenerative conditions such as Alzheimer’s disease, with vasomotion being strongly linked to the function of the glymphatic system^19,20^. However, in these studies it could be argued that they are not investigating ‘vasomotion’ as defined in this study, but rather NVC at low frequencies loosely centred around 0.1Hz. Further studies are needed in both anaesthetised and awake mice where vasomotion is stimulated (for example by low blood pressure as in this study or the inhibition of Nitric Oxide Synthase^21^ to confirm whether vasomotion in the mouse has the same structure as that seen in the rat sensory cortex. Utilising non-invasive neural measurements, such as GCAMP in combination with simultaneous hemodynamic measurements would allow further characterisation of vasomotion and how it interacts with ongoing spontaneous NVC, both in rats and mice.

### Vasomotion as a travelling wave

In this study, we have shown (Fig 4) that vasomotion has the capacity to manifest as a robust coherent travelling wave aligned with the geometry of the middle cerebral artery. This has potential importance for spontaneous connectivity research especially those that have shown the appearance of travelling waves at similar frequencies ^22–24^. If the wave shown in this study had its origin bilaterally at the circle of Willis, it is likely that homologous connected regions may be found across both cerebral hemispheres. Therefore, a more robust analysis of connectivity data is potentially needed, especially in human cohorts with disease where vasomotion may be an emergent property of a pathological cerebral circulation^25^.

### Source of the LFP vasomotion signal

From this study, we do not know the precise cause of the LFP vasomotion signal, which was observed when large vasomotion was present in the absence of burst-suppression neuronal activity. There are likely two potential mechanisms. The first would be a physical movement artefact caused by the hemodynamic vasomotion oscillation. However, if this were the case it would be expected that all electrode channels would exhibit the same signal. Generally, not all channels in the electrode showed this (see Fig 4 and 5). The second possibility is that the electrode is picking up electrical activity from the vessel wall when an artery is close to a recording site. The phase relationship of 0.43 seconds aligns with studies assessing conducted vasodilation in isolated arterial vessel preparations^26^ which showed calcium waves moving along the endothelial cell layer causing subsequent vessel dilation. An intriguing possibility from our results is the potential that electrodes can be designed with sites of varying impedance so that some sites measure neuronal activity where others can record vascular-based responses. This is an area of future investigation.

### Limitations and future research

This study was performed in anaesthetised rats, which may limit the generalisation of the results.^27^. Whilst the anaesthetised preparation has limitations particularly in terms of potential confounds in NVC mechanisms; it does allow for greater control and the elimination of equally large confounds found in the awake preparation such as from locomotion^28^. In this study, we took advantage of the fact urethane anaesthesia causes a drop in blood pressure, which is known to enhance cerebral vasomotion^11^. In future combined studies of both awake and anaesthetised preparations, where appropriate will add further knowledge to understanding cerebral vasomotion and its relevance to human cerebrovascular health as both a biomarker of pathology and potential therapeutic strategy.

## Methods

### Animal preparation and surgery

Surgical and experimental procedures were performed in accordance with the Animal (Scientific procedures) Act 1986, with approval from the UK home office. Eighteen Female *Hooded Lister rats* were kept at 22°C, in a 12-hour dark/light cycle, with food and water supplied *ad libitum*. Rats weighing between 250 to 400g were anaesthetised with Urethane at 1.25g/kg i.p. with additional doses of 0.1ml administered if required. Atropine was administered at 0.4mg/kg s.c. to lessen mucous secretions during surgery. Following anaesthesia core temperature was maintained at 37°C throughout surgery and experimental procedures using a homoeothermic heating blanket system with rectal temperature monitoring (Harvard Instruments). Animals were tracheotomised allowing for artificial ventilation and continuous monitoring of end-tidal CO_2_ (CapStar-100, CWE systems, USA). Blood gas measurements were taken and along with the CO_2_ monitoring were used to adjust the ventilator parameters to keep the animal within physiological limits. Cannulation of the femoral arteries was performed to monitor mean arterial blood pressure (MABP) and to allow blood samples to be taken for blood gas measurement. A femoral vein was cannulated to maintain MABP between 100-110 mmHg through the infusion of phenylephrine at 0.13-0.26 mg/h^29,30^. Subjects were placed in a stereotaxic frame (Kopf Instruments). The skull surface was exposed and the area of bone overlying the right somatosensory and motor cortices was thinned to translucency, using a dental drill, with saline cooling the surface. A plastic well (20mm diameter) with saline input and output ports was fixed over the right somatosensory cortex using dental cement. The well was constantly infused with saline to increase transparency of the window and to reduce optical specularities from the skull.

### Placement of whisker stimulating electrodes

Stainless steel electrodes, with a 2mm uninsulated tip were subcutaneously inserted into the left whisker pad between rows A/B and C/D. Electrical pulses were applied to the electrodes to confirm stimulation of the majority of the whisker pad. Stimuli produced no changes in MABP, pCO_2_ or heart rate. This suggests the stimulation produced no changes in systemic physiology which could affect cortical activity or hemodynamic changes.

### Two-Dimensional Optical Imaging Spectroscopy (2D-OIS)

Cortical haemodynamic changes were measured using 2D-OIS. The recording of haemodynamic changes using 2D-OIS rather than BOLD fMRI has several advantages for the study of NVC. The principal advantage is the ease with which 2D-OIS can be used with multi-channel / multi-probe electrophysiological recording with minimal electrical interference, allowing for concurrent measures of neural activity. Additional advantages include the more comprehensive haemodynamic measures that 2D-OIS offers over BOLD fMRI, with relative measures of total haemoglobin and saturation. Lastly 2D-OIS generally has better spatial and temporal resolutions than high field BOLD fMRI and thus can detect smaller magnitude haemodynamic changes elicited by smaller changes in neural firing. The region of thinned skull overlying the right somatosensory cortex was illuminated in sequence with four wavelengths of light, using a Lambda DG-4 high speed galvanometer-based filter changer with integrated light source (Sutter Instrument Company, Novato, California, USA). A CCD camera (1M30P, Teledyne DALSA Inc, US) operating at 4x4 binning recorded images of the cortical surface, with each image pixel representing 75x75µm of the cortical surface. The camera had a quantum efficiency of 28% at 500nm. The camera captured images at 32Hz and was used to synchronise filter changing for each of the four wavelength exposures, giving an effective frame rate of 8Hz. The four wavelengths were chosen specifically as two pairs (495 nm ± 31 FWHM and 559 nm ± 16 FWHM; 575 nm ± 14 FWHM and 587 nm ± 9 FWHM). The wavelengths in each pair were chosen to have similar total absorption coefficients, thus sampling the same tissue volume, but to have specific absorption coefficients for oxyhaemoglobin (HbO_2_) and deoxyhaemoglobin (Hbr) that were as different as possible to maximise the signal to noise ratio. Recorded images were subject to spectral analysis using a path length scaling algorithm (PLSA), which was made from a modified Beer-Lambert law with a path length correction factor, for more detail see ^31,32^. The algorithm required a value for the baseline concentration of haemoglobin in tissue, which was set at 100 µM with saturation set at 80% for arterial regions based on previous measurements ^33^. 2D-OIS produced spatio-temporal ‘images’ of cortical haemodynamics with measures of HbO_2_, Hbr and total haemoglobin (Hbt). Hbt was the aspect of haemodynamics chosen for subsequent analyses of hemodynamic changes. Each hemodynamic trial represents a Micromolar change in concentration from its own pre-stimulus baseline period.

### Localisation of the whisker barrel and surrounding regions for placement of recording electrodes

2D-OIS was initially used to functionally localise the whisker barrel and motor cortex areas within the thinned cranial window (See Figure 1), by detecting the cortical hemodynamic changes occurring in responses to brief stimulation of the whisker pad. The stimulation paradigm consisted of 30 stimulation trials with an inter-stimulus-interval of 25s, with 300µs pulse width stimuli presented for 2s at 5Hz with a constant current of 0.8mA. Individual stimulus presentation trials were trial averaged before being subject to spectral analysis. A general linear model (GLM) was applied to the data and used to select a spatial region showing the greatest stimulus related increase in Hbt (Figure 1c). In a first group of animals (*N*=12) two regions corresponding to whisker sensory and whisker motor cortex were identified in each animal, and this enabled accurate placement of two multi-channel electrode arrays into each region. In a second group of animals (*N*=6) a single electrode was placed in whisker sensory region in combination with an oxygen electrode (OxyLite Pro, Oxford Optronix UK) to measure baseline levels of oxygenation. Care was taken to avoid surface vasculature at the sites of electrode placement. A very small hole was drilled in the thinned skull directly over the selected regions. The Dura was pierced with a 27-gauge needle and the 16-channel silicone linear array electrodes (100 µm spacing, area of each site 177 µm^2^, Neuronexus Technologies) were inserted normal to the cortical surface to a depth of 1600 µm, using a stereotaxic arm (Kopf instruments). The probes were coupled to a pre-amplifier which was in turn connected to a data acquisition unit via fibre optic cable (Medusa Bioamp, Tucker Davis Technologies (TDT), Florida). Stimulus-evoked and spontaneously occurring neural recordings were sampled at 24.41 KHz with a 16-bit resolution, with the data collected using openEX software (Tucker Davis Technologies, Florida) which also handled trigger timings and data storage.

### Experimental paradigm

The experimental design took advantage of the fact that urethane anaesthesia causes a decrease in baseline blood pressure. It is also known that vasomotion oscillations are increased when blood pressure is low. Therefore, the experimental design was to perform a 2100s experiment measuring baseline hemodynamic (2D-OIS) and neural activity(TDT) in which a constant infusion of phenylephrine (0.13-0.26 mg/h) was started after 290s. This method in our previous publications^34–37^ established an extremely stable baseline for neurovascular experiments with the absence of vasomotion. In this experiment we investigated how changing the magnitude of cerebral vasomotion effects ongoing baseline neurovascular relationships. A representative animal and experimental design are shown in Figure 2. In this representative animal that has electrodes placed in right whisker and motor regions (Figure 2c and d), baseline blood pressure is low prior to the infusion of phenylephrine (at 290s) and then increases to normal physiological values at approximately 1000s (Figure 2g).

### Pre-post phenylephrine time series analysis (Arterial Hbt, MABP and Oxygen tension)

Data were compared using a paired t-test. The Shapiro-Wilk test of Normality was used to assess normality. If Shapiro-Wilk test result was significant, then Wilcoxon Signed Ranks Test was used as an alternative to compare datasets. Results of statistical tests were considered significant if *p*<0.05.

### Creation of averaged vasomotion trials for spatiotemporal 2D-OIS and neural analysis

To understand how vasomotion is structured across the cortex we developed a methodology to assess its spatial specificity and assess whether it was driven by neuronal activity. In the period prior to MABP increase we selected Hbt peaks of vasomotion, in most animals we were able to identify 50-75 identifiable peaks. For each peak, 10s of spatial data was extracted before and after the peak for Hbt data. Each dataset represented a vasomotion trial (similar to trials extracted following whisker stimulation) and all the trials were then averaged for each electrode site used. As we collected simultaneous 2D-OIS data we could also extract both the LFP and MUA data aligned to the spontaneous Hbt peaks.

For a further analysis (Figure 6) we sub-sorted the animals into two further classes. The first, where the animals had vasomotion in the absence of burst suppression prior to phenylephrine infusion (11 datasets from 8 animals with three having both whisker and motor electrodes included). The second, after phenylephrine infusion that showed strong burst suppression with little or no remaining vasomotion (11 datasets from 6 animals with 4 having both whisker and motor electrodes included). In a subset of 10 animals, we also performed 2s electrical whisker stimulation (as described above) after electrode insertion and MABP stabilisation.

### Assessment of spontaneous NVC relationships using a Kernel analysis

The Hbt data was bandpass filtered (0.01–0.5 Hz) and deconvolved with gamma band LFP (30-80 Hz) using a kernel-based approach ^38^. The relationship is expressed as:

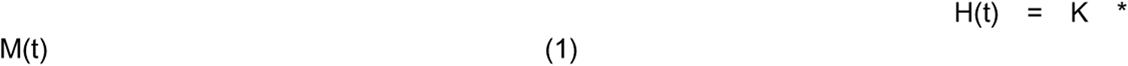

where H(t) is the predicted Hbt, *M(t)* is the LFP, ∗ denotes the convolution operator, and *K* is the kernel. The predicted Hbt was calculated by convolving the kernel and measured LFP. The proportion of Hbt that can be explained by neural activity was evaluated based on the correlation between measured and predicted Hbt across conditions.

## Acknowledgements

This work was funded by Medical Research Council UK grant [Grant number MR/M013553/1].

For the purpose of open access, the author has applied a Creative Commons Attribution (CC BY) licence to any Author Accepted Manuscript version arising.

## Code, Data, and Materials Availability

Data and code presented in this paper are available on the ORDA Repository: https://figshare.com/s/23820d820274b41e9ec6

## Author contributions

RW: Data analysis, Manuscript preparation PP: Data Collection, Experiment design

LB: Data Collection, Experiment design, Technology Design MO: Data analysis, Manuscript preparation and editing.

CH: Data analysis, Manuscript preparation

JB: Experiment design, Data analysis, Manuscript preparation and editing.

## References

1 Logothetis, N. K. & Wandell, B. A. Interpreting the BOLD signal. Annual review of physiology 66, 735–769 (2004). 10.1146/annurev.physiol.66.082602.092845

2 Friston, K. J. Functional and effective connectivity: a review. Brain Connect 1, 13–36 (2011). 10.1089/brain.2011.0008

3 van den Heuvel, M. P. & Hulshoff Pol, H. E. Exploring the brain network: a review on resting-state fMRI functional connectivity. Eur Neuropsychopharmacol 20, 519–534 (2010). 10.1016/j.euroneuro.2010.03.008

4 Zlokovic, B. V. Neurodegeneration and the neurovascular unit. Nature medicine 16, 1370–1371 (2010). 10.1038/nm1210-1370

5 Roy, C. S. & Sherrington, C. S. On the Regulation of the Blood-supply of the Brain. The Journal of physiology 11, 85–158 117 (1890). 10.1113/jphysiol.1890.sp000321

6 Mayhew, J. E. et al. Cerebral vasomotion: a 0.1-Hz oscillation in reflected light imaging of neural activity. NeuroImage 4, 183–193 (1996). 10.1006/nimg.1996.0069

7 Goldman, D. & Popel, A. S. A computational study of the effect of vasomotion on oxygen transport from capillary networks. J Theor Biol 209, 189–199 (2001). 10.1006/jtbi.2000.2254

8 Mateo, C., Knutsen, P. M., Tsai, P. S., Shih, A. Y. & Kleinfeld, D. Entrainment of Arteriole Vasomotor Fluctuations by Neural Activity Is a Basis of Blood-Oxygenation-Level-Dependent "Resting-State" Connectivity. Neuron 96, 936–948 e933 (2017). 10.1016/j.neuron.2017.10.012

9 van Veluw, S. J. et al. Vasomotion as a Driving Force for Paravascular Clearance in the Awake Mouse Brain. Neuron 105, 549–561 e545 (2020). 10.1016/j.neuron.2019.10.033

10 Biswal, B. B. & Hudetz, A. G. Synchronous oscillations in cerebrocortical capillary red blood cell velocity after nitric oxide synthase inhibition. Microvasc Res 52, 1–12 (1996). 10.1006/mvre.1996.0039

11 Hudetz, A. G., Biswal, B. B., Shen, H., Lauer, K. K. & Kampine, J. P. Spontaneous fluctuations in cerebral oxygen supply. An introduction. Adv Exp Med Biol 454, 551–559 (1998).

12 Clement, E. A. et al. Cyclic and sleep-like spontaneous alternations of brain state under urethane anaesthesia. PloS one 3, e2004 (2008). 10.1371/journal.pone.0002004

13 Bouskela, E. Vasomotion frequency and amplitude related to intraluminal pressure and temperature in the wing of the intact, unanesthetized bat. Microvasc Res 37, 339–351 (1989). 10.1016/0026-2862(89)90051-4

14 Bouskela, E. & Grampp, W. Spontaneous vasomotion in hamster cheek pouch arterioles in varying experimental conditions. Am J Physiol 262, H478–485 (1992). 10.1152/ajpheart.1992.262.2.H478

15 Collin, O., Zupp, J. L. & Setchell, B. P. Testicular vasomotion in different mammals. Asian J Androl 2, 297–300 (2000).

16 Ma, Y. et al. Wide-field optical mapping of neural activity and brain haemodynamics: considerations and novel approaches. Philosophical transactions of the Royal Society of London. Series B, Biological sciences 371 (2016). 10.1098/rstb.2015.0360

17 Kozberg, M. G. et al. Loss of spontaneous vasomotion precedes impaired cerebrovascular reactivity and microbleeds in a mouse model of cerebral amyloid angiopathy. bioRxiv (2024). 10.1101/2024.04.26.591414

18 Munting, L. P. et al. Spontaneous vasomotion propagates along pial arterioles in the awake mouse brain like stimulus-evoked vascular reactivity. Journal of cerebral blood flow and metabolism : official journal of the International Society of Cerebral Blood Flow and Metabolism 43, 1752–1763 (2023). 10.1177/0271678X231152550

19 Harrison, I. F. et al. Impaired glymphatic function and clearance of tau in an Alzheimer’s disease model. Brain 143, 2576–2593 (2020). 10.1093/brain/awaa179

20 Smith, A. J. & Verkman, A. S. The "glymphatic" mechanism for solute clearance in Alzheimer’s disease: game changer or unproven speculation? FASEB J 32, 543–551 (2018). 10.1096/fj.201700999

21 Lindauer, U., Megow, D., Matsuda, H. & Dirnagl, U. Nitric oxide: a modulator, but not a mediator, of neurovascular coupling in rat somatosensory cortex. Am J Physiol 277, H799–811 (1999). 10.1152/ajpheart.1999.277.2.H799

22 Majeed, W., Magnuson, M. & Keilholz, S. D. Spatiotemporal dynamics of low frequency fluctuations in BOLD fMRI of the rat. Journal of magnetic resonance imaging : JMRI 30, 384–393 (2009). 10.1002/jmri.21848

23 Thompson, G. J. et al. Neural correlates of time-varying functional connectivity in the rat. NeuroImage 83, 826–836 (2013). 10.1016/j.neuroimage.2013.07.036

24 Thompson, G. J., Pan, W. J., Magnuson, M. E., Jaeger, D. & Keilholz, S. D. Quasi-periodic patterns (QPP): large-scale dynamics in resting state fMRI that correlate with local infraslow electrical activity. NeuroImage 84, 1018–1031 (2014). 10.1016/j.neuroimage.2013.09.029

25 Rayshubskiy, A. et al. Direct, intraoperative observation of ∼0.1 Hz hemodynamic oscillations in awake human cortex: implications for fMRI. NeuroImage 87, 323–331 (2014). 10.1016/j.neuroimage.2013.10.044

26 Uhrenholt, T. R., Domeier, T. L. & Segal, S. S. Propagation of calcium waves along endothelium of hamster feed arteries. American journal of physiology. Heart and circulatory physiology 292, H1634–1640 (2007). 10.1152/ajpheart.00605.2006

27 Gao, Y. R. et al. Time to wake up: Studying neurovascular coupling and brain-wide circuit function in the un-anesthetized animal. NeuroImage 153, 382–398 (2017). 10.1016/j.neuroimage.2016.11.069

28 Eyre, B. et al. The effects of locomotion on sensory-evoked haemodynamic responses in the cortex of awake mice. bioRxiv, 2021.2012.2016.472794 (2021). 10.1101/2021.12.16.472794

29 Golanov, E. V., Yamamoto, S. & Reis, D. J. Spontaneous waves of cerebral blood flow associated with a pattern of electrocortical activity. Am J Physiol 266, R204–214 (1994). 10.1152/ajpregu.1994.266.1.R204

30 Nakai, M. & Maeda, M. Scopolamine-sensitive and resistant components of increase in cerebral cortical blood flow elicited by periaqueductal gray matter of rats. Neuroscience letters 270, 173–176 (1999). 10.1016/s0304-3940(99)00491-7

31 Berwick, J. et al. Fine Detail of Neurovascular Coupling Revealed by Spatiotemporal Analysis of the Hemodynamic Response to Single Whisker Stimulation in Rat Barrel Cortex. J Neurophysiol 99, 787–798 (2008). 10.1152/jn.00658.2007

32 Berwick, J. et al. Neurovascular coupling investigated with two-dimensional optical imaging spectroscopy in rat whisker barrel cortex. European Journal of Neuroscience 22, 1655 (2005).

33 Kennerley, A. et al. Concurrent fMRI and optical measures for the investigation of the hemodynamic response function. Magnetic Resonance in Medicine 54 (2005).

34 Berwick, J. et al. Fine detail of neurovascular coupling revealed by spatiotemporal analysis of the hemodynamic response to single whisker stimulation in rat barrel cortex. Journal of neurophysiology 99, 787–798 (2008). 10.1152/jn.00658.2007

35 Boorman, L. et al. Long-latency reductions in gamma power predict hemodynamic changes that underlie the negative BOLD signal. The Journal of neuroscience : the official journal of the Society for Neuroscience 35, 4641–4656 (2015). 10.1523/JNEUROSCI.2339-14.2015

36 Boorman, L. et al. Negative blood oxygen level dependence in the rat: a model for investigating the role of suppression in neurovascular coupling. The Journal of neuroscience : the official journal of the Society for Neuroscience 30, 4285-4294 (2010). 10.1523/JNEUROSCI.6063-09.2010

37 Boorman, L. W. et al. Bidirectional alterations in brain temperature profoundly modulate spatiotemporal neurovascular responses in-vivo. Communications Biology 6, 185 (2023). 10.1038/s42003-023-04542-6

38 Aster, R. C., Borchers, B., & Thurber, C. H. . Parameter Estimation and Inverse Problems Third Edition, 211–233 (2019). 10.1016/B978-0-12-804651-7.00013-4

